# Low-Dimensional Representations of Visuomotor Coordination in Natural Behavior

**DOI:** 10.1101/2024.03.30.587357

**Authors:** Ashima Keshava, Maximilian A. Wächter, Franca Boße, Thomas Schüler, Peter König

## Abstract

Understanding how the eyes, head, and hands coordinate in natural contexts is a critical challenge in visuomotor coordination research, often constrained by sedentary tasks, cued actions, or restricted settings. To address this gap, we conducted an experiment where participants could self-generate pick-and-place actions on a life-size shelf in a virtual environment, recording concurrent gaze and body movements. Subjects exhibited intricate translation and rotation movements of the eyes, head, and hands during the task. We employed a time-dependent principal component analysis to study the relationship between the movements of the eyes, head, and hands relative to the onset of the action. We reduced the overall dimensionality into 2D representations, capturing up to 65% of the movement variance just in time with the actions. Our analysis revealed a synergistic coupling of the eye-head and eye-hand systems, as well as a strong coupling within the head-hand system. The eyes synchronized with the head and hands close to the action onset, with variations in coupling observed in horizontal and vertical planes, indicating distinct mechanisms for coordination in the brain. Crucially, the head and hands remained tightly coupled throughout the observation period, suggesting a shared neural code driving these effectors. Notably, the low-dimensional representations of the eye-head-hand movement vectors showed maximum predictive accuracy of the action’s location ~200ms before the action onset, highlighting just-in-time coordination among the three effectors. Furthermore, the predictive accuracy was significantly influenced by the location of the upcoming action. Our study emphasizes the differential visuomotor control subject to the task structure, providing insights into the dynamic interplay of eye, head, and hand movements during natural behavior.

**NEW & NOTEWORTHY:** Studying natural, self-initiated, complex visuomotor coordination, we observe low-dimensional dynamics with distinct patterns along horizontal and vertical axes. The eye’s horizontal movement showed notable independence, aligning with head and hand movements just in time for action. Notably, around critical events, the dimensionality of the complex movements is further reduced, indicating dynamic correspondence of eye-head-hand coordination.

## INTRODUCTION

Eye-hand coordination is a defining characteristic of everyday activities. Routine tasks involving object interactions require coordination between multiple sensorimotor systems to execute bodily transformations and affect environmental changes. Studies of tasks like sandwich-making [1] or tea-making [2] show how the visual information is sampled by the eye to direct and guide hand movements to accomplish goals iteratively until the task is adequately terminated. In this sense, eye movements serve a predominant function of planning and assisting manual interactions in the real world. Hence, visuomotor coordination is necessary to sample relevant visual information to output appropriate motor responses.

Task constraints, available sensory information, and the cognitive context can affect eye-hand coordination [3, 1, 4]. Studies have shown that gaze targets earmark upcoming sensorimotor events [5, 6]. There is also consistent evidence that gaze control supports the planning of hand movements in object manipulation [7]. These findings have shown eye movements gather information from the environment proactively and in anticipation of upcoming manual action [8, 9]. Vision-for-action in naturalistic tasks can be accomplished with just-in-time representations [10] where gaze fixations relay information for an action right before the action. While eye-hand coordination has been studied in various task contexts, there are still gaps in the literature concerning how the structure of the external world might affect these coordination strategies.

Studies in visuomotor control have been investigated in both head-restrained and unrestrained contexts. In a head-restrained setup, stimuli are primarily shown on a computer screen in a limited space [7, 11, 12, 13]. It is known that eye-in-head signal accuracy declines at larger angles in head-fixed experiment designs[14] and raises concerns about the generalizability of results from sedentary experimental setups to real-world human behavior. In more naturalistic contexts with head unrestrained setups, subjects are shown targets on a computer screen [15, 16], in a limited horizontal space [17, 18] such as a table or countertop and often constrained to only one dimension (often in the horizontal plane) even when participants were allowed to stand [19]. Experimenters often cue movements in naturalistic and head unrestrained contexts, and subjects do not make self-generated, proactive movements. Furthermore, the cued movements are frequently practiced beforehand. Subjects usually sit during the experiments and have little freedom to move the head and trunk outside the experimental setup’s confines. These studies have undoubtedly enhanced our understanding of how the eye, head, and hands coordinate in naturalistic contexts. Yet, they do not fully capture the nature of unrestrained and self-generated movements.

Recent theories of embodied cognition have emphasized the importance of studying self-generated, goal-directed movements as a foundation for understanding sensorimotor processes [20, 21]. Naturalistic approaches offer a vital addition to traditional, highly controlled laboratory studies [22], allowing researchers to investigate behavior in contexts that more closely mirror everyday experience. Several influential studies have pioneered this direction by enabling participants to engage in self-initiated strategizing and unconstrained interactions with real-world objects [2, 23, 24, 25, 13]. While these studies provide foundational insights, their analyses tend to be primarily descriptive, examining eye, head, and hand movements in isolation without a formal model of their joint coordination. In addition, most rely on manual annotation of gaze and action from video, making the methods difficult to scale or generalize across participants. Hence, there are also methodological concerns about how we study visuomotor coordination.

In this study, we ask: How do the eyes, head, and hands coordinate during natural, self-initiated movements, and how is this coordination influenced by the spatial structure of the task space? Advances in virtual reality (VR) now allow researchers to explore such questions in controlled yet immersive settings, using wearable eye and body trackers to record kinematic signals with high precision [26, 27, 28, 29]. Here, we used VR to capture rich, three-dimensional (3D) data streams from the eyes, head, and hands as participants performed naturalistic sorting tasks. Unlike earlier studies, our approach uses automated preprocessing and computational modeling to uncover how the eye-head-hand effectors coordinate. Our approach allows us to scale from descriptive accounts to a more formal, computational ethology perspective [30] of visuomotor coordination.

Natural behavior is highly dynamic and complex, especially when multiple effectors operate in 3D space. Understanding coordination mechanisms thus requires methods that can reduce this complexity while preserving its latent structure. To this end, we used low-dimensional representations to capture the shared variance of eye-head-hand movements. High-dimensional kinematic data, reflecting translations and rotations of various effectors, were projected into a latent subspace using Principal Component Analysis (PCA), which identifies patterns of coordinated variation across signals. This low-dimensional representation provides a compact summary of visuomotor behavior and helps characterize the set of states the body occupies during natural action [31, 32, 33]. This approach enables a principled analysis of complex, high-dimensional behavior, offering insights into the underlying structure and dynamics of sensorimotor coordination.

We explored human volunteers’ movement trajectories while sorting objects on a 2m wide and 2m high life-size shelf in VR. They performed pick-and-place actions iteratively until a cued goal was achieved. Crucially, the participants were free to move and generate their own movements, unconstrained by time, allowing them to act as naturally as possible in the virtual environment. The tasks were generic (involving pick-and-place actions) but novel (requiring planning to sort the objects according to varying rules), making the findings relevant to visuomotor coordination in a proactive and natural context. We focussed on how eye-head-hand coordination unfolded in this context by studying the movements in a low-dimensional subspace and cross-validating how these representations predict the timing and location of upcoming action events. By addressing these questions, our study extends previous work on natural behavior through a formal and generalizable framework that captures the dynamics of visuomotor coordination.

## MATERIALS & METHODS

### Ethics Statement

The Ethics Committee of the University of Osnabrück approved the study (Ethik-37/2019). Before each experimental session, subjects gave their informed consent in writing. They also filled out a questionnaire regarding their medical history to ascertain they did not suffer from any disorder/impairments that could affect them in the virtual environment. After obtaining their informed consent, we briefed them on the experimental setup and task.

### Participants

A total of 55 participants (39 females, mean age = 23.9 ± 4.6 years) were recruited from the University of Osnabrück and the University of Applied Sciences Osnabrück. Participants had normal or corrected-to-normal vision and no history of neurological or psychological impairments. They either received a monetary reward of C7.50 or one participation credit per hour.

### Apparatus & Procedure

For the experiment, we used an HTC Vive Pro Eye head-mounted display (HMD)(110° field of view, 90Hz, resolution 1080 × 1200 px per eye) with a built-in Tobii eye-tracker with 120 Hz sampling rate. With their right hand, participants used an HTC Vive controller to manipulate the objects during the experiment. The HTC Vive Lighthouse tracking system provided positional and rotational tracking and was calibrated for a 4m x 4m space. We used the 5-point calibration function provided by the manufacturer to calibrate the gaze parameters. To ensure the calibration error was less than 1°, we performed a 5-point validation after each calibration. Due to the study design, which allowed many natural body movements, the eye tracker was calibrated repeatedly during the experiment after every 3 trials. Furthermore, subjects were fitted with HTC Vive trackers on both ankles, both elbows and one on the midriff. The body trackers were also calibrated subsequently to give a reliable pose estimation using inverse kinematics in the virtual environment. We designed the experiment using the Unity3D v2019.4.4f1 and SteamVR game engine and controlled the eye-tracking data recording using HTC VIVE Eye-Tracking SDK SRanipal v1.1.0.1.

The experiment consisted of 16 objects placed on a shelf with a 5×5 grid. The objects were differentiated based on two features: color and shape. Each object could be differentiated based on one of four high-contrast colors (red, blue, green, and yellow) and four 3D shapes (cube, sphere, pyramid, and cylinder). The objects had an average height of 20cm and a width of 20cm. The shelf was designed with a height and width of 2m with five rows and columns of equal height, width, and depth. Participants were presented with a display board on the right side of the shelf where the trial instructions were displayed. Subjects were also presented with a red buzzer to end the trial once they finished the task. The physical dimensions of the setup are illustrated in Figure 1A. The shelves’ horizontal eccentricity from the shelf’s center extended to 89.2 cm in the left and right directions. Similarly, the vertical eccentricity of the shelf extended to 89.2 cm in the up-down direction from the center-most point of the shelf. This ensured that the task setup was symmetrical in both the horizontal and the vertical directions. The objects on the top-most shelves were placed at a height of 190cm and on the bottom-most shelves at 13cm from the ground level.

**Figure 1:**
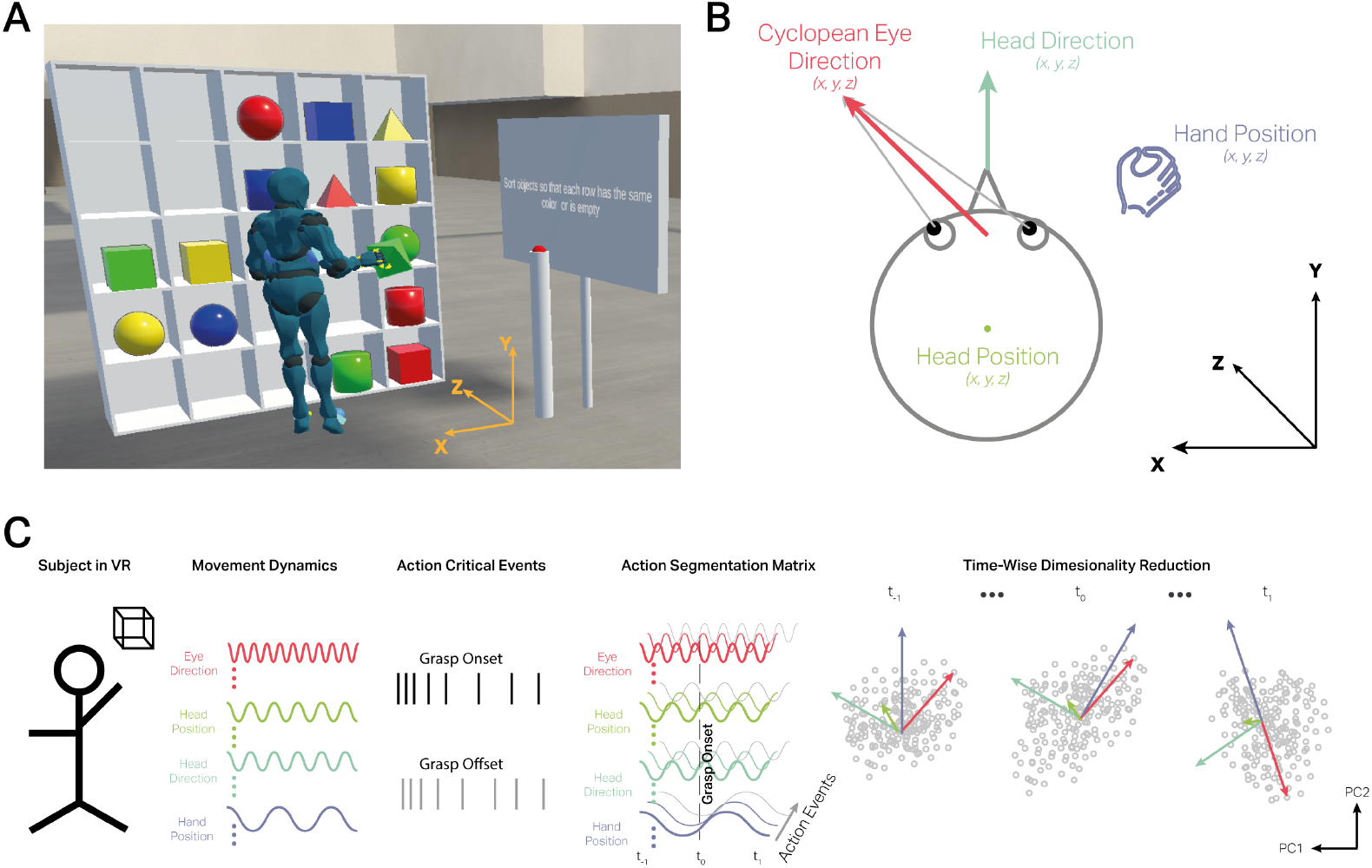
Experimental Setup. **A**. In a virtual environment, 27 participants sorted 16 different objects based on two features (color or shape) while we measured their eye and body movements. The objects were randomly presented on a 5×5 shelf at the beginning of each trial and were cued to sort objects by shape and/or color. All participants performed 24 trials in total with no time limit. **B**. Positions of the eye, head, and hand are recorded in 3D Cartesian coordinates. For subsequent analyses, eye direction and hand position are expressed in the head-centered frame (i.e., relative to the head origin and axes). **C**. To understand the coupling of the different sources of data from the head, eye, and hand in different reference frames, we performed dimensionality reduction analysis on the measured signals relative to action critical events of grasp onset (when the hand touches the object to pick up and the VR controller trigger is pressed) and grasp offset (when controller trigger is released and the hand drops the object on the desired shelf after object displacement). For each object displacement, we segmented the data to 1s before and after the action events. For each subject, the data matrix was composed of *G* × *M* × *T*, where *G* denotes object displacements, *M* denotes the 12 features (four data streams in *x, y, z* coordinates), and *T* denotes each time point relative to the action events. We applied Principal Component Analysis at each time point to the *G* × *M* matrix and reduced its dimensionality to *G* × 2. The resulting 2D representation across time would then elucidate the contribution of the original data sources to the principal components and their explained variance.

### Experimental Task

Subjects performed two practice trials where they familiarized themselves with the handheld VR controller and the experimental setup. In these practice trials, they were free to explore the virtual environment and displace the objects. After the practice trials, subjects were asked to sort objects based on one and/or two features of the object. Each subject performed 24 trials in total, with each trial instruction (as listed below) randomly presented twice throughout the experiment. The experimental setup is illustrated in Figure 1A. The trial instructions were as follows:

1. Sort objects so that each row has the same shape or is empty
2. Sort objects so that each row has all unique shapes or is empty
3. Sort objects so that each row has the same color or is empty
4. Sort objects so that each row has all unique colors or is empty
5. Sort objects so that each column has the same shape or is empty
6. Sort objects so that each column has all unique shapes or is empty
7. Sort objects so that each column has the same color or is empty
8. Sort objects so that each column has all unique colors or is empty
9. Sort objects so that each row has all the unique colors and all the unique shapes once
10. Sort objects so that each column has all the unique colors and all the unique shapes once
11. Sort objects so that each row and column has each of the four colors once.
12. Sort objects so that each row and column has each of the four shapes once.

### Data pre-processing

We recorded the 3D data representing the position and direction of the eye, head and hand in global 3D coordinates. The eye direction and the hand position vectors were subsequently expressed in the head-centered reference frame by applying a rigid-body transformation (translation and rotation) relative to the head origin and orientation. By orientation, we mean the yaw or pitch of the head. Also, the distinction between head and cyclopean eye positions arises from a fixed anatomical offset between the two. The eye-tracking software models the cyclopean eye as a point fixed in the head but offset from the head origin due to its anatomical location. Furthermore, the software outputs the direction vectors as quaternions in UnityVR. For easier understanding, we transformed these vectors from quaternions to 3D unit vectors. So, a forward-pointing vector with 0° rotation along the horizontal and vertical axes is represented as [0, 0, 1]. This ensures that orientation (yaw and pitch) is expressed as a 3D unit vector in Cartesian space rather than spherical angles. Figure 1B illustrates the vector representations of the eye, head, and hand position and direction vectors.

At the outset, we downsampled the data to 40 Hz at the outset so all samples were equally spaced. The following sections explain the steps we took to process the raw data and arrive at the magnitude of translation and rotation movements made by the eye, head, and hand.

### Gaze data

Using the 3D gaze direction vector for the cyclopean eye, we calculated the orientation of the eye in Euler angle in the horizontal *θ*_h_ and vertical *θ*_v_ directions. Vector samples were sorted by their timestamps. Each sample’s 3D gaze direction vector is represented in (*x, y, z*) coordinates as a unit vector that defines the direction of the gaze in VR world space coordinates. In VR, the *x* coordinate corresponds to the left-right direction, *y* in the up-down direction, *z* in the forward-backward direction. We computed the horizontal (*θ*_h_) (eq. 1) and the vertical (*θ*_v_) (eq. 2) angles in degrees. The formula calculates the 3D gaze direction vector projection onto the x-z (horizontal) and y-z (vertical) planes w.r.t the *z* axis. In Python, we used the *atan*2 function, which correctly interprets the sign of both components to give the full range of angles in 360°.

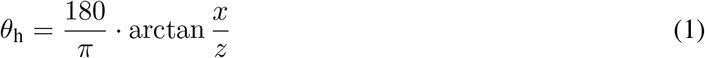

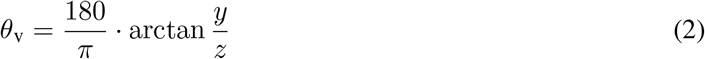

The computed horizontal and vertical angles were used to depict the distribution of gaze direction during the experiment and were not used in further analysis.

### HMD data

From the HMD we obtained the global head position vector and the head direction vector. Using the 3D head direction vector, we calculated the horizontal (*ρ*_h_, eq. 3) and vertical (*ρ*_v_, eq. 4) projection angles onto the x-z (horizontal) and y-z (vertical) planes respectively. The resulting angles provided the head’s horizontal (yaw) and vertical (pitch) directions in the VR world space across time. These angles were similarly used to depict the distribution of the head direction movements during the experiment and were not used in further analysis.

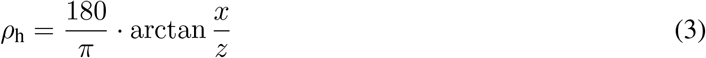

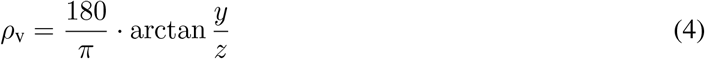

At the beginning of each trial, subjects were asked to stand still facing the shelf for 3s at a set location. Thus, we could estimate the initial position of the head from the average position of the HMD in the 3s period. Subsequently, we calculated the deviations of the HMD from the initial position to arrive at the magnitude and direction of translation movements. For each time point after the initial 3s period, we subtracted the initial position from the current position in 3D coordinates. Hence, we could estimate the degree of head translation movements in the left-right and up-down directions.

### VR hand controller data

Subjects used the trigger button of the HTC Vive controller to virtually grasp the objects on the shelf and displace them to other locations. In the data, the trigger was recorded as a boolean, which was set to TRUE when subjects pressed the trigger button on the hand controller to initiate an object displacement and was reset to FALSE when the trigger button was released to place the object on the shelf. Using the position of the controller in the world space, we determined the locations from the shelf where a grasp was initiated and ended. We also removed trials where the controller data showed implausible locations in the 3D space. These faulty data were attributed to the loss of tracking during the experiment. Next, we disregarded grasping periods where the objects’ origin and final displacement locations were the same on the shelf.

Next, we calculated the angular position of the hand with respect to the head positions in each trial. As described above, the x-coordinate corresponds to the left-right direction, y in the up-down direction, z in the forward-backward direction. Using the 3D Cartesian coordinates (*x, y, z*) of the controller position (*Hand*_(*x,y,z*)_), HMD position (*Head*_(*x,y,z*)_) and HMD direction unit vector 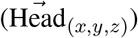 in world space, we calculated the horizontal and the vertical angular position of the hand with respect to the head. The horizontal (*ϕ*_h_) and vertical (*ϕ*_h_) angular position of the hand was calculated as follows:

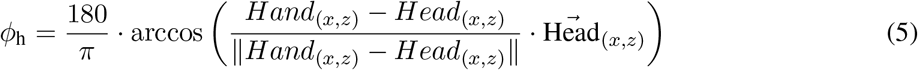

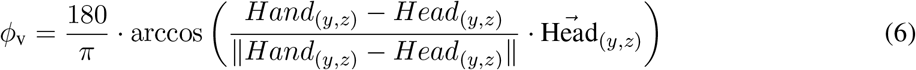

where ∥∥ denotes the norm of the vector, and (·) denotes the dot product. The computed angles were used to get the distribution of the horizontal and vertical positions of the hand w.r.t the head position.

To summarize, we calculated the projection angles in the horizontal and vertical planes of the eye, head, and hand to depict these effectors’ overall movement during the experiment.

### Data Analysis

After pre-processing, we were left with data from 27 subjects with 554 trials in total and *Mean* = 20.51, *SD* = ±2.20 trials per subject. Furthermore, we had eye-tracking data corresponding to 5664 grasping actions with 12.85, *SD* = ±1.91 object displacements per trial and subject.

### Principal Components Analysis

Given the naturalistic setting of our experimental setup, with complex movements of the head, eye, and hand, we wanted to understand the contributions and coupling of each of these effectors while performing object pickup (grasp onset) and dropoff (grasp offset) actions. We first epoched the data using the grasp onset and offset triggers. We selected a time window of 1s before and after the trigger. Thus, each epoch consisted of *x, y, z* coordinates of the head position, head direction, eye direction relative to head direction, and hand position relative to the head position at each time point spaced 0.025s apart. Each subject’s feature matrix consisted of a 3D matrix of 80 time points with 12 features for each grasping event. We further explored the time-wise low-dimensional representation of this matrix using Principal Component Analysis.

For each subject, the input matrix for PCA was composed of *G* × *M* × *T*, where *G* denotes the grasping epoch, *M* denotes the 12 features (four data streams in x, y, z coordinates), and *T* denotes each time point relative to the action events. For each time point in T, we standardized *G* × *M* matrix to zero mean and unit standard deviation. This was done to make sure each feature contributed equally to the PCA. We then applied PCA to the *G* × *M* matrix at each time point in *T*. We used the ‘sklearn.preprocessing.PCA’ python package to compute the eigenvalues and principal components. We then used the ‘transform()’ function to reduce the dimensionality of the matrix from *G* × *M* to *G* × 2. Thus, for each subject, we obtained *T* × 2 eigenvalues and 2 × *M* × *T* coefficients of the eigenvectors. The eigenvalues were used to show the explained variance of the principal components (PCs) across time. The coefficients provided the loadings of each *M* feature onto the PCs. Hence, for each subject, we could ascertain the evolution of the contribution of the features in the PCA subspace across time.

### Cosine Similarity of Factor Loadings

To understand the contributions of the individual features to the PCA subspace, we calculated the directional similarity of the variables. We computed the cosine similarity of the *x* and *y* coordinates in the PCA subspace using eq 7 for each eye-head, eye-hand, and head-hand pair.

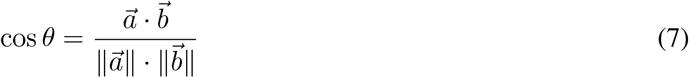

Using the values of cos *θ*, we could determine if the coefficients of the eigenvector were aligned in the same direction (cos *θ* = 1), opposite directions (cos *θ* = −1) or orthogonal (cos *θ* = 0) across the different time points. Hence, through the cosine similarity analysis, we could ascertain how the *x* and *y* components of the eye-head, eye-hand, and head-hand pairs correlated across time.

### Generalization & Prediction

To explore the generalizability of the PCA subspace across subjects. We used *N*-fold cross-validation, where *N* corresponds to the number of subjects. For each fold, we divided the dataset into train and test sets, where data from *N* − 1 subjects was used for training the model, and the left-out dataset from 1 subject was used for testing it. During training, we applied PCA to the *A* × *M* matrix at each time point of the grasping epochs, where *A* denotes all grasping epochs across *N* − 1 subjects. After applying PCA, we obtained the *A* × 2 reduced matrix in PCA subspace. Using the *A* × 2 matrix, we trained a kernel-based support vector machine (SVM) to predict the location of the action at each time point. We used the sklearn.svm.SVC python package for training and prediction. We used the default parameters for the model and did not perform any hyper-parameter optimization. During testing, we standardized the *G × M* matrix of the left-out subject, applied the PCA weights obtained from the training set to reduce its dimensionality to *G* × 2, and recorded the mean prediction accuracy of the trained model on the left-out data. This analysis was repeated for the grasp offset events. In this manner, we could test the generalizability and predictive power of the PCA subspace across time for grasp onset and offset events.

### Sources of Variance in Prediction

We further explored the sources of variance in the predictive accuracy and the influence of the action location. We used a linear model to test the hypothesis of whether the ground truth row and column location of the grasp onset and offset events influenced the peak predictive accuracy of the low-dimensional space. This helped us infer the robustness of the low-dimensional representations with respect to the action locations on the shelf.

We modeled the relationship between the peak predictive accuracy dependent on the ground truth row and column location of the action. The row and columns were denoted as ordinal variables from 1-5. For the rows, numbers 1 to 5 represent the top-to-bottom rows. For the columns, 1-5 represented left to right columns. The model fit was performed using restricted maximum likelihood (REML) estimation [34] using the lme4 package (v1.1-26) in R 3.6.1. We used the bobyqa optimizer to find the best fit using 20000 iterations. Using the Satterthwaite method [35], we approximated degrees of freedom of the fixed effects. The modeling was done separately for grasp onset and offset events. The model in Wilkinson notation [36] is denoted as follows:

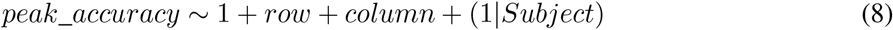

## RESULTS

Twenty-seven healthy human volunteers performed object-sorting tasks in VR on a 2m high and 2m wide life-size shelf. Participants performed 24 trials each. In each trial, participants sorted 16 randomly presented objects on the shelf based on a cued task and iteratively performed pick and place actions until the cued sorting was achieved. Figure 1A illustrates the experimental setup in VR. The experimental task consisted of sorting objects by their features. Each object was differentiated by color and shape. As the objects were randomly presented to the participants, they made reaching movements to different locations in space, such as reaching for objects closer to their feet or to the top of their heads. With multiple sorting tasks, we ensured that they manipulated objects from multiple locations on the shelf. We analyzed each grasping action and its corresponding location irrespective of the overall sorting task. Such a setup guarantees that subjects make different movements across the height and breadth of the shelf, and we can capture the global attributes of coordinated translation and rotation movements of the eye, head, and hand. Hence, we recorded ecologically valid and proactive movements within a large spatial context.

For this study, we utilized the 3D coordinates of four continuous data streams. Namely, the head position of the participants, head direction, i.e., the unit direction in which the head was oriented, the cyclopean eye (average of left and right eye) unit direction within the head, and the hand angular position relative to the head position. The direction vectors described the rotation of movement by indicating how the eye and head were oriented or moving in the 3D space. While the head direction vector described the head’s orientation in world space, the eye direction vector denoted the eye’s orientation in the head reference frame. These 3D movement vectors were represented in (*x, y, z*) coordinates (Figure 1B). We analyzed data from 27 participants, 5664 grasp actions with 209.77 ± 3.41 grasps per subject.

At the outset, we down-sampled the data to 40 Hz so that the continuous samples were equally spaced at 25ms. The down-sampling also smoothed the movement trajectories. We segmented the data streams relative to the onset of a grasp. We chose the time window from 1s before the grasp onset to 1s after. The time window was selected from the study of grasp timing in previous literature [2, 37, 11]. We similarly epoched the data relative to the grasp offset with the same window lengths. This allowed us to capture the eye, head, and hand movement trajectories while reaching the object, grasping it, and guiding it to a desired location. For both grasp onset and grasp offset events, the continuous data comprised 80 time-points starting from −1s to 1s relative to the action event, where time point 0 marked the onset or offset of said action. We then used PCA to reduce the overall dimensionality of the segmented data at each time point. The original data comprised of 12 dimensions (four data streams, each with 3d coordinates). We performed a time-wise PCA on each of the 80 time points to illustrate how the low-dimensional representation of the ongoing visuomotor coordination morphed relative to an action event and how the individual eye, head, and hand orientations contributed to these low-dimensional representations across time. Figure 1C depicts the data segmentation steps and PCA analysis approach.

### Complexity of Natural Behavior

Participants performed complex movements, which led to varied translation and rotation of the head. To understand the range of movements made by the participants, we transformed the position and direction vectors into projection angles on the x-z (horizontal) and y-z (vertical) planes. Figure 2A shows the joint distribution of the initial position of the participants at the beginning of each trial in the horizontal and vertical planes. All participants started the trial from a fixed position in VR. The mean initial position of the participants in the horizontal plane was 0.0*m, SD* = ± 0.03, and in the vertical plane was 1.63*m* ± 0.07. The variance in the vertical direction corresponds to the variance in the participants’ height. From the initial positions, we calculated the translation-based deviations of the participants during the trials. Figure 2B shows the bi-variate distribution of the horizontal and vertical translation head movements from the initial position. In the horizontal plane, the mean deviation of the head was −0.04*m* ± 0.34; in the vertical plane, it was −0.07*m* ± 0.14. As evidenced by the spread of the distributions, participants made more translation movements in the horizontal plane. The downward movements show the translation of the head needed to interact with objects in the lower shelf locations. Figure 2C shows the bi-variate distribution of head direction vector projection angles in the horizontal and vertical planes. The mean rotation in the horizontal plane was −2.80° ± 18.90, and −15.84° ± 18.70 in the vertical plane. The data shows participants made symmetrical horizontal head rotation movements. However, they tended to direct their heads downward in the vertical plane. Taken together, the head movement vectors spanned the dimensions of the shelf. The distributions of the translation and rotation of the head showed no outlying behavior of the participants.

**Figure 2:**
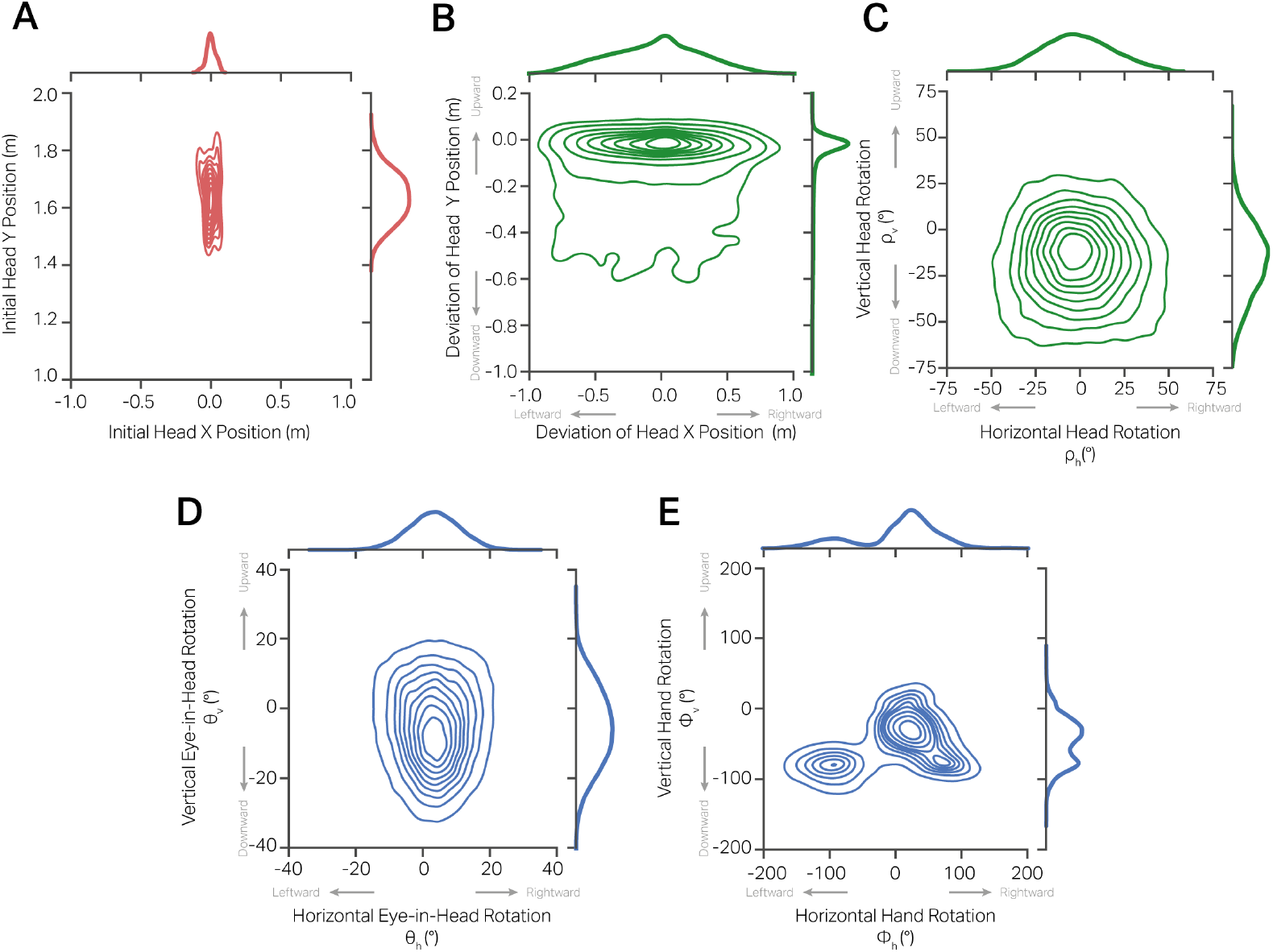
Complexity of natural movements. Plots show joint and marginal distributions of the participants’ head, eye, and hand movements in the horizontal and vertical planes. **A**. Bi-variate distribution of head position at the start of each trial. **B**. Bi-variate distribution of the deviations of head movements from the initial position in the trials in the horizontal (abscissa) and vertical (ordinate) directions. **C**. Joint distribution of projection angles of head direction vector during the trials. When participants aligned their heads straight ahead, facing the shelf, and along the z-axis, the rotation in the horizontal and vertical direction was 0°. **D**. Joint distribution of the projection angles of eye direction in the head reference frame in the horizontal and vertical planes during the trials. When participants aligned their eyes and head straight ahead, facing the shelf, and along the z-axis, the rotation in the horizontal and vertical direction was 0°. **E**. Bi-variate distribution of hand position in the head reference frame.

We were interested in the range of motion exhibited by the participants’ eye and hand movements during the trials. Figure 2D shows the bi-variate distribution of the eye rotation in the head reference frame. The mean eye projection angle in the horizontal plane was 3.42°*SD* = ±7.50. Similarly, the mean projection angle in the vertical plane was −5.49° ± 10.98. Figure 2E shows the distribution of the projection angles of the hand position in the head reference frame. The mean angle in the horizontal plane was 4.15°± 61.43; in the vertical plane, the mean angle was −46.12° ± 31.79. The data shows that the eye had symmetric rotations in both axes, with a more significant variation in the vertical plane. Moreover, the average eye direction was slightly off-center with respect to the head direction, i.e., within the head, the eye was slightly rightward and downward. This is likely an artifact of the task, as subjects were instructed to use their right hand to manipulate the objects.

The above data shows the complexity of natural behavior in a naturalistic spatial context. As the eye, head, and hand vectors show a complex structure, we could not isolate the individual horizontal and vertical components of these vectors and directly correlate them. Hence, to decipher the associations between the eye, head, and hand movements, we chose to study their relationship in a low-dimensional plane.

### Low Dimensional Representations of Visuomotor Coordination

Due to the complex range of motion exhibited by the eye, head, and hand, we studied the properties of the different movement vectors not as isolated systems but together. When the eye, head, and hand movement vectors are coordinated, we would find redundancies in the whole system, and fewer vectors could explain the joint coordination. As a first step, to understand the dynamic contribution of the different vectors relative to the start and end of a grasp, we performed a time-wise Principle Component Analysis (PCA). For each of these action-critical events, we performed the PCA using 3D vectors of the head position, head direction, eye direction, and hand position. We applied time-wise PCA to data from each subject. In doing so, we could ascertain first the average variance explained by the low-dimensional representations of the movement vectors across time and, subsequently, the contributions of the position and direction vectors to these low-dimensional representations.

### Grasp onset

To understand the low-dimensional representation of eye, head, and hand movement vectors relative to grasp onset events, we performed time-wise PCA on head position, head direction, eye direction, and hand direction 3D vectors (*x, y, z*) totaling 12 features per subject. Figure 3A shows the explained variance ratio (eigenvalues) of the principal components (PCs) across time, from 1s before action onset to 1s after. The explained variance of the top two PCs increases as the action onset approaches and subsequently reduces. At 1s before the action onset, the mean explained variance ratio for PC1 is 0.31(*SD* ± = 0.03), and of PC2 is 0.18 ± 0.01. At time point 0, at the onset of grasp when the hand makes contact with the object to be picked up, PC1 and PC2 have a mean eigenvalue of 0.42 ± 0.03 and 0.22 ± 0.02, respectively. At the end of the grasp window, at 1s after the grasp onset, PC1 and PC2 have a mean eigenvalue of 0.33 ± 0.04 and 0.18 ± 0.01, respectively. Notably, the peak explained variance ratio is achieved slightly before the action onset at −0.19*s*, 95%*CI* = [−0.24, −0.14]. Throughout the grasp epoch, the other PCs exhibit an explained variance ratio of less than 0.15. This analysis shows that the 12 movement features can be reduced to a lower dimensional space consisting of two PCs that together explain more than 50% of the variance across time and are most informative near the action onset.

**Figure 3:**
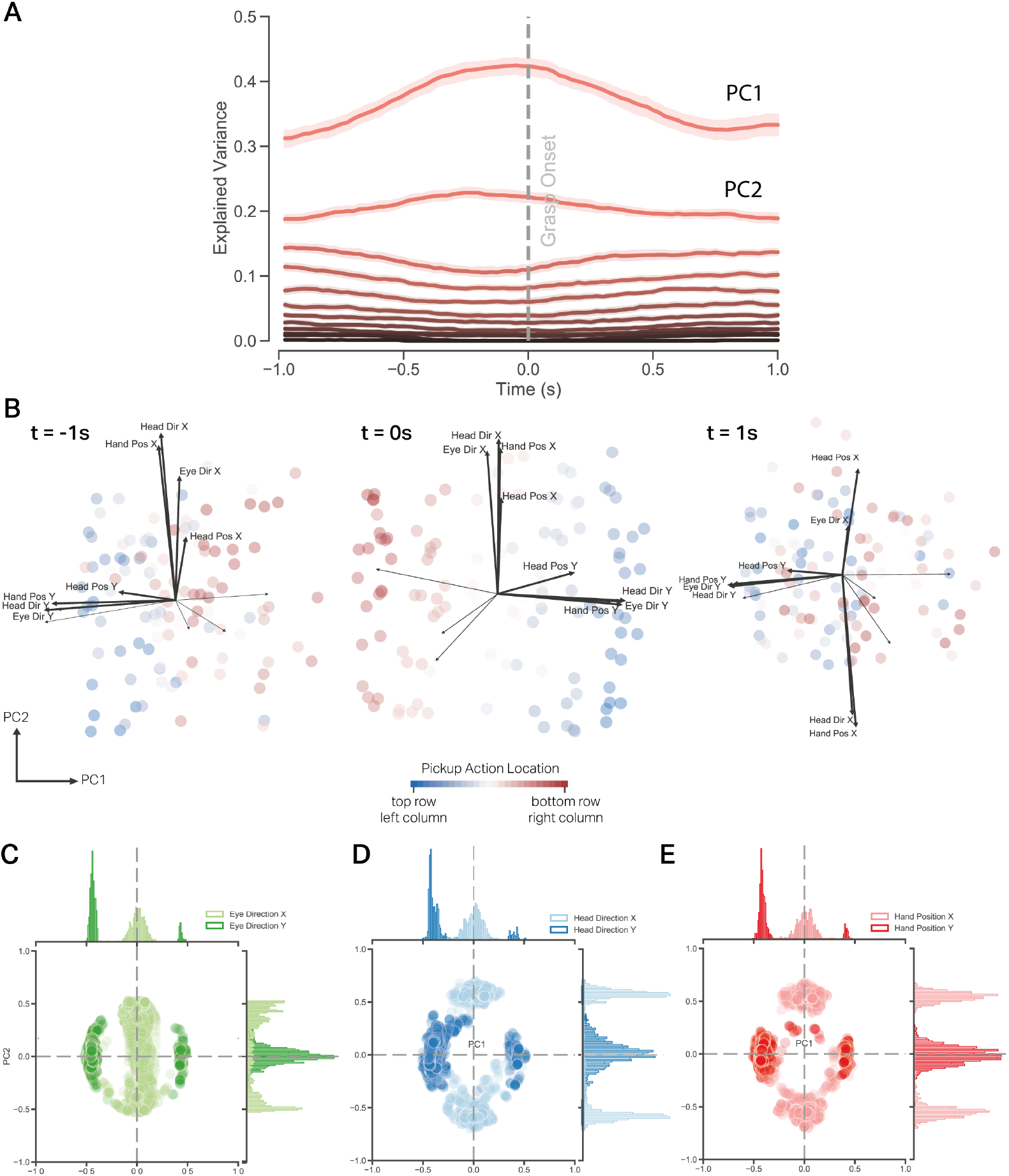
PCA analysis of 3D eye direction, head direction, head position, and hand position vectors. **A**. Temporal profile of eigenvalues of the 12 principle components (PCs) 1s before and after the grasp onset at time 0s. The solid lines denote the mean variance explained over the 27 participants by each PC, and the shaded region depicts the 95% confidence interval. **B**. Projection of exemplar subject data onto the first two PCs for time points −1s (left), 0s (middle), 1s (right). Each dot within a time point denotes a pickup action projected onto PC1 and PC2, with the action colored according to the spatial location of the pickup action on the 5×5 shelf. The black arrow vectors represent the loadings of the original feature variables to the first two PCs. The magnitude of the vector denotes the strength of the correlation between the original variable and the two PCs. The direction provides the sign of the correlation (positive or negative). **C**. Bivariate distribution of the contributions of the horizontal and vertical eye direction vectors on the two PCs across the different time points and their respective marginal distributions. **D**. The joint contribution of the horizontal and vertical head direction vectors on the two PCs. **E**. The contribution of the horizontal and vertical hand position vectors on the two PCs across the different time points. As seen from these distributions, the horizontal components of the eye, head, and hand contributed primarily to PC2, and the vertical components contributed to PC1.

Next, to further understand the structure of the low-dimensional space, we plotted the contribution of each movement vector to this low-dimensional space. Figure 3B shows the low dimensional projection of the data from an exemplar subject onto PC1 and PC2 for time points −1s before grasp onset, 0s at grasp onset, and 1s after grasp onset. The direction of the component loadings illustrates its correlation with the first two PCs, and the length of the vector shows the magnitude of the correlation. The data are further colored according to the location where the upcoming pick-up action is performed. In this low dimensional representation, we see that 1s before the grasp onset, the data begin to cluster according to the shelf height where the pickup action will subsequently happen. Furthermore, the loadings of the vertical (*y*) components of the head direction, eye direction, and hand position are similar and have a small angular deviation. The *x* components of these vectors have significant angular deviation and contribute differentially to the two PCs. At grasp onset (time=0s), the data are well clustered according to the pickup action location, and the horizontal and vertical components of the head direction, eye direction, and hand position point in the same direction. At time 1s after the grasp onset, the data are not clustered according to the pickup action location, and horizontal and vertical components of the movement vectors ‘disengage.’ The evolution of the PCA subspace across the entire time window is shown in Supplementary Material Movie 1. This analysis of the loadings of individual features shows their evolution within the low-dimensional space described by PC1 and PC2. Moreover, at grasp onset, the horizontal and vertical components of the movement vectors are orthogonally represented, point in the same direction at grasp onset, and convey the same information to the two principal components.

Each principal component is composed of a linear combination of the original input variables. The weights of the original variables in the low-dimensional space indicate their respective correlations to the two PCs. The sign of the weights denotes the direction of the relationship with the PCs. To understand the overall contribution of the original movement vectors across all time points and participants, we charted their joint contributions to the 2D eigenvectors (PC1, PC2) as shown in Figure 3C, D, E. The marginal distributions of the original variables in the principal component space show the vertical components of the eye, head direction, and hand position consistently contribute to the PC1, and the horizontal components correlate with the PC2. As seen from the plots, the horizontal components have a bimodal distribution on PC2 where equal proportions of participants’ data contributed similarly to PC2 but in opposite directions at different points in time. Table 1 details the central tendency of the absolute value of the feature loadings in the low-dimensional space. The PC1 and PC2 consisted of primarily vertical and horizontal components of the movement vectors, respectively, where the vertical components accounted for the largest proportion of the variance across time.

**Table 1:** Mean absolute factor loadings of the original variables on the first two principal components across the grasping epoch.

| variable | PC1 | PC2 |
| --- | --- | --- |
| Head Direction X | 0.06 (0.05) | 0.56 (0.07) |
| Head Direction Y | 0.41 (0.03) | 0.07 (0.05) |
| Head Direction Z | 0.11 (0.08) | 0.13 (0.10) |
| Head Position X | 0.07 (0.05) | 0.25 (0.12) |
| Head Position Y | 0.29 (0.08) | 0.08 (0.06) |
| Head Position Z | 0.16 (0.10) | 0.14 (0.10) |
| Eye Direction X | 0.05 (0.04) | 0.38 (0.13) |
| Eye Direction Y | 0.43 (0.02) | 0.06 (0.05) |
| Eye Direction Z | 0.36 (0.07) | 0.09 (0.08) |
| Hand Position X | 0.06 (0.06) | 0.54 (0.07) |
| Hand Position Y | 0.40 (0.03) | 0.05 (0.04) |
| Hand Position Z | 0.38 (0.04) | 0.10 (0.08) |

### Grasp offset

We repeated the above analysis for the object grasp offset events, i.e., when the object in hand is placed on the desired shelf. This was considered an action-critical event as the eye, head, and hand would have to coordinate to guide the object to a set location and could be meaningfully different from the coordination required for reach movements. Figure 4A shows the explained variance ratio of twelve PCs when decomposing the head position, head direction, eye direction, and hand position vectors in 3D. Here again, the explained variance ratio of the first two PCs is around 0.50 throughout the time period and 0.60 close to the grasp offset. At timepoint −1s before the grasp offset, PC1 and PC2 had a mean explained variance ratio of 0.34 ± 0.03 and 0.18 ± 0.01. At time point 0, the mean explained variance ratio for PC1 was 0.39 ± 0.03 and 0.19 ± 0.01, respectively. Finally, at time point 1s after the grasp offset, PC1 and PC2 exhibited a mean explained variance ratio of 0.32 ± 0.03 and 0.19 ± 0.02, respectively. Similar to the grasp onset event, the increased explained variance ratio for the first two PCs indicates a higher correlation between the eye, head, and hand vectors. Nonetheless, the explained variance ratio of the first two PCs sums to 0.58 compared to the grasp onset events, which is at 0.64. This lower explained variance ratio at the grasp offset events suggests a lower correlation between the movement vectors.

**Figure 4:**
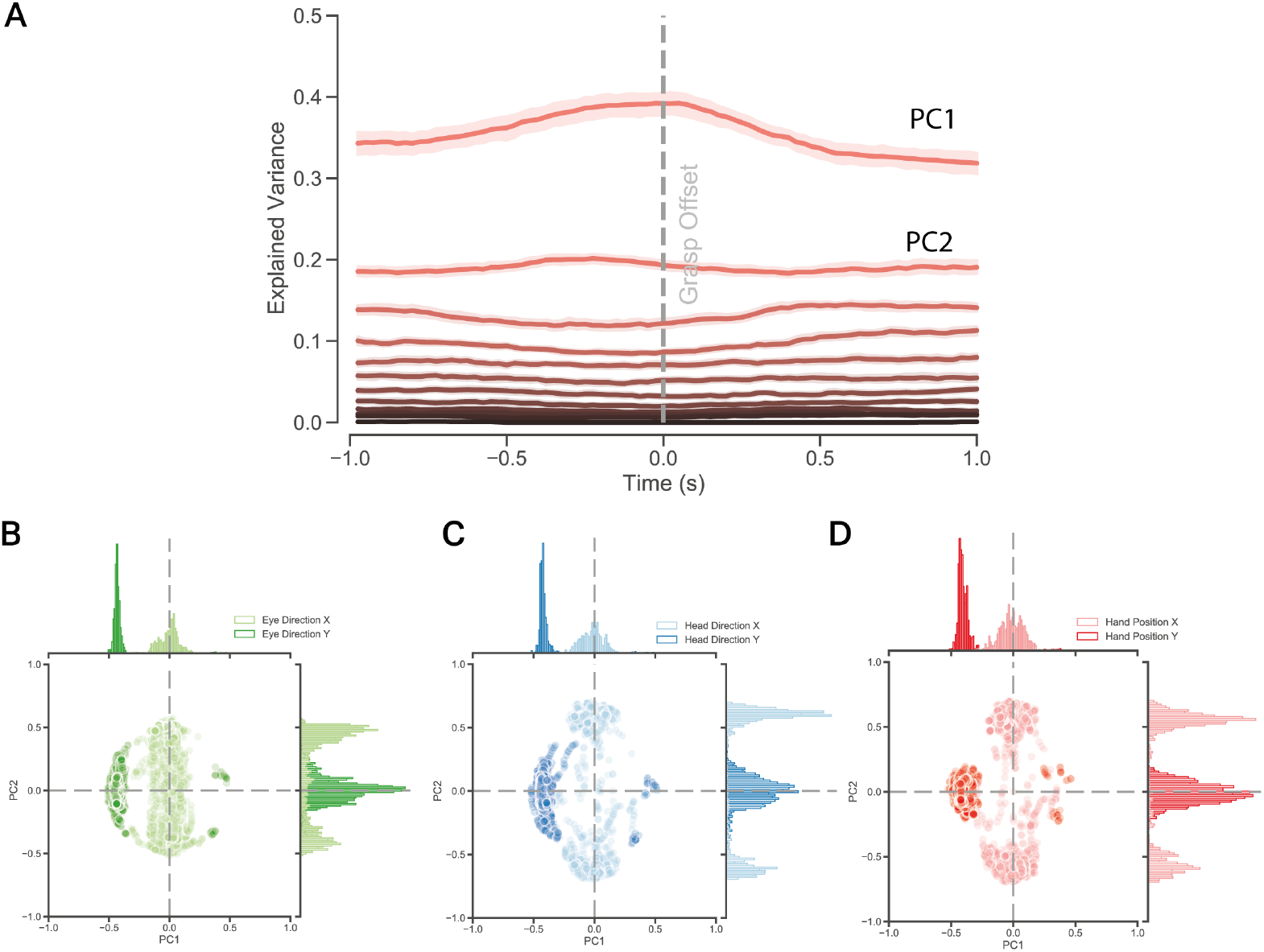
PCA analysis of 3D eye direction, head direction, head position, and hand position vectors. **A**. Temporal profile of eigenvalues of the 12 principal components (PCs) 1s before and after the grasp onset at time 0s. The solid lines denote the mean explained variance ratio over the 27 participants by each PC, and the shaded region depicts the 95% confidence interval. **B**. Bivariate distribution of the contributions of the horizontal and vertical eye direction vectors on first two PCs across the time points and participants, and their respective marginal distributions. **C**. The joint contribution of the horizontal and vertical head direction vectors on the two PCs. **E**. The contribution of the horizontal and vertical hand position vectors on the two PCs across the different time points. As seen from these distributions, the horizontal components of the eye, head, and hand contributed primarily to PC2, and the vertical components contributed to PC1.

As before, we checked for consistency of the contributions of the different movement vectors to the low-dimensional space. As seen from the joint distribution of the loadings of the original variable (Figure 4B, C, D), the horizontal components of the movement vectors consistently contributed to PC2, and the vertical components to PC1 across different time points and participants. Table 2 details the mean and standard deviation of the absolute contribution of each variable on the first two PCs. In sum, the low-dimensional space primarily captured the correlations of the horizontal and vertical components of the movement vectors, where the vertical components explained more variance in the data than the horizontal components.

**Table 2:** Mean factor loadings of the original variables on the first two principal components during grasp offset.

| variable | PC1 | PC2 |
| --- | --- | --- |
| Head Direction X | 0.07 (0.06) | 0.57 (0.11) |
| Head Direction Y | 0.42 (0.03) | 0.08 (0.07) |
| Head Direction Z | 0.12 (0.08) | 0.14 (0.11) |
| Head Position X | 0.06 (0.05) | 0.19 (0.12) |
| Head Position Y | 0.30 (0.08) | 0.08 (0.07) |
| Head Position Z | 0.17 (0.10) | 0.14 (0.10) |
| Eye Direction X | 0.06 (0.05) | 0.36 (0.13) |
| Eye Direction Y | 0.43 (0.02) | 0.07 (0.06) |
| Eye Direction Z | 0.34 (0.09) | 0.13 (0.12) |
| Hand Position X | 0.07 (0.06) | 0.50 (0.11) |
| Hand Position Y | 0.40 (0.03) | 0.06 (0.04) |
| Hand Position Z | 0.39 (0.04) | 0.08 (0.07) |

### Cosine Similarity Analysis

In the above section, we performed time-wise PCA on action-critical events of grasp onset and grasp offset using head position, head direction, eye direction, and hand position 3D vectors. The results showed a larger explained variance of the first two PCs, where this ratio was maximum just before the action events. PCA provides a subspace where data variance is maximized across the principal components. Using a vector similarity analysis, we can further explore relationships between the original variables within this subspace and identify which variables contribute similarly to the explained variance. Hence, exploration of the subspace can lead to a deeper understanding of the data structure, such as revealing groups of variables that might be part of the same underlying process or phenomenon.

To quantify the source of the increase in the explained variance ratio close to the action onset and offset events, we used cosine similarity analysis. Cosine similarity measures the correlation between the factor loadings in the 2D PCA subspace. PCA aims to identify patterns of similarity and differences across variables by transforming them into PCs based on their covariance. Hence, using cosine similarity to analyze the correlation of variables in this transformed space further complements the goals of PCA. It helps understand the structure and relationships between variables beyond mere dimensionality reduction. When the PC loadings point in the same direction, they indicate a high positive correlation with each other. Similarly, when they point in opposite directions, they indicate a high negative correlation. When they are orthogonal to each other, they indicate no correlation at all. Hence, using cosine similarity, we could ascertain the evolution of the correlation between the horizontal and vector components of the movement vectors before and after the action-critical events.

In grasp onset epochs, we calculated the pairwise 6cosine similarity between the *x* (horizontal) and *y* (vertical) components of the eye direction, head direction, and hand position 3D vector for each time point across subjects. Figure 5A shows the mean cosine similarity of the horizontal and vertical components of the head and eye direction vectors and the standard deviation. At time point 0, the horizontal and vertical components had a mean similarity of 0.99 ± 0.002 and 0.99 ± 0.001, respectively. Between eye and hand factor loadings (Figure 5B), we observed a mean similarity of 0.99 ± 0.002 in the horizontal direction and 0.99 ± 0.001 in the vertical direction at grasp onset. Between head and hand factor loadings (Figure 5C), we observed a remarkable consistency throughout the action epoch where the similarity of the horizontal components was 0.99 ± 0.0003 and for vertical components 0.99 ± 0.001 at time 0s. Taken together, the vertical components of the eye, head, and hand factor loadings were well correlated throughout the grasp onset epoch. However, the horizontal components were correlated around 0.5s before the grasp onset, and this correlation reduced drastically shortly after the grasp event was triggered. Throughout the time course, the hand and head vectors varied in the same direction, both in the horizontal and the vertical planes, and exhibited a strong correlation.

**Figure 5:**
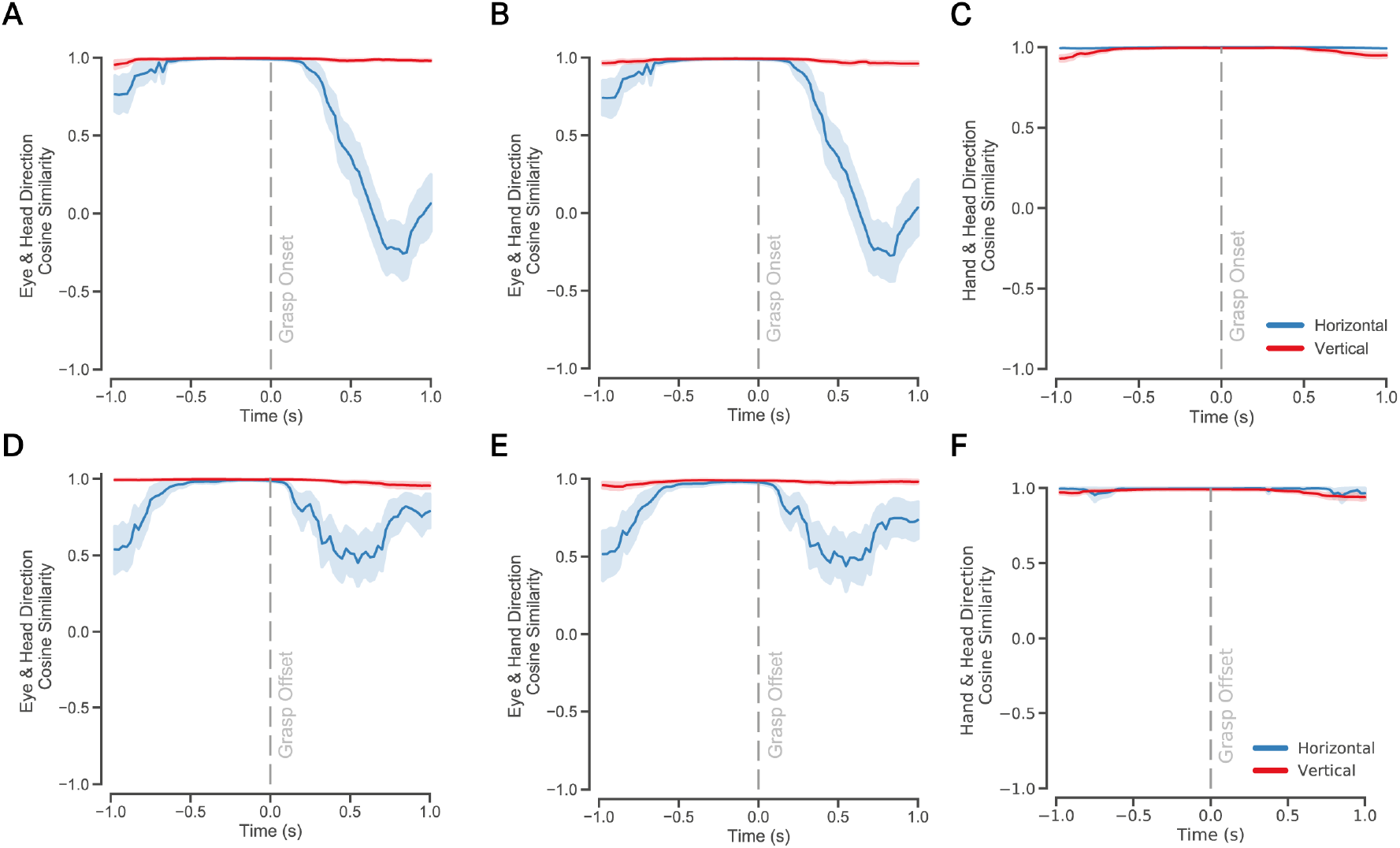
Cosine similarity between head, eye, and hand direction vectors for grasp onset and grasp offset events. **A**. Cosine similarity between the horizontal and vertical components of head and eye direction 3D vectors during grasp onset events. The solid lines denote the mean cosine similarity across 27 participants, and the shaded region denotes the standard error or mean. A cosine similarity value of 1 indicates a perfect correlation between the vectors in the PCA subspace, whereas a value of −1 denotes a perfect negative correlation. A value of 0 indicates the vectors are not related to each other. **B**. Cosine similarity between the horizontal and vertical components of eye direction and hand position 3D vectors during grasp onset events. **C**. Cosine similarity between the horizontal and vertical components of head and eye direction vectors during grasp offset events. **D**. Cosine similarity between the horizontal and vertical components of eye direction and hand position vectors during grasp offset events.

We repeated the above analysis for the grasp offset events. First, we calculated the pairwise cosine similarity between the horizontal and vertical components of the eye, head, and hand factor loadings in the PCA subspace. Figure 5D illustrates the average similarity of eye-head loadings over subjects across the different time points. At time point 0, the average cosine similarity was 0.98 ± 0.004 in the horizontal direction and 0.95 ± 0.01 in the vertical direction. Between eye-hand factor loadings (Figure 5E), the horizontal and vertical components were perfectly aligned at time 0s and showed an average similarity of 0.97 ± 0.008 and 0.98 ± 0.003, respectively. Between head-hand factor loadings (Figure 5E), we observed a striking similarity as before, where the horizontal components had a mean similarity measure of 1.00 ± 0.003 and 0.99 ± 0.003 for the vertical components. Taken together, there was again a strong coupling between the vertical components of the eye, head, and hand. In contrast, the horizontal components were aligned in the same direction briefly before the grasp offset. Further, the head direction and the hand position vectors covaried in the same direction and showed very high correlations.

In sum, exploring the PCA subspace with the loadings of the original variables showed varied aspects of visuomotor coordination. Namely, the vertical components of the eye, head, and hand vectors were almost perfectly aligned in the low-dimensional space, suggesting a near perfect correlation. The horizontal component of the eye direction vector, on the other hand, was only briefly oriented in the same direction as the head direction and hand position vectors at about 0.5s before the grasp onset and offset. This window of alignment of the vectors also coincides with the increase in the explained variance ratio of the first two PCs before the action onset. Crucially, the head direction and the hand position vectors tracked in complete unison throughout the time window. Thus, the similarity analysis of the effectors in the PCA subspace showed distinct coordination mechanisms for the horizontal and vertical components of the visuomotor system.

### Generalization and Predictive Accuracy of the Low-dimensional Space

To further expound on the generalizability and predictive power of these low-dimensional structures, we predicted the location of the action at each time point based on the PCA-transformed data. For each time point *t*, we pooled the data from *N* − 1 subjects and standardized it to zero mean and unit standard deviation. We then reduced the dimensionality of the data at each time point into a 2D space. For each time point, we trained a kernel-based support vector machine (SVM) to classify the location of the upcoming action. We used leave-one-subject cross-validation to arrive at the validated test accuracy. For each time point, we standardized the test data and transformed it using the PCA weights from the training data. We then computed the prediction accuracy of the trained SVM on the test data at that time-point. We performed the above steps until each of the 27 subjects’ data was used as test data for all time points ranging from 1s before and after the grasp onset. We repeated this analysis for the grasp offset events as well. Our analysis provided an aggregated prediction accuracy of the PCA-transformed training and test datasets. Thus, using cross-validation, we could generalize the information encapsulated in the PCA subspace and ensure the prediction accuracy was not affected by the peculiarities of single subjects.

The prediction of the object pickup action location across the grasp onset epoch provided a greater understanding of the evolution of the PCA subspace as seen in Figure 6A. At the 1s before grasp onset, the prediction accuracy on the test data is low at *Mean* = 0.20, 95%*CI* = [0.17, 0.23]. At time 0, the accuracy increases to 0.62 [0.52, 0.72]. At 1s after the grasp onset, the prediction accuracy is reduced to 0.12 [0.10, 0.13]. This shows that the PCA subspace encodes more relevant information about the upcoming action close to the action onset. The maximum predictive accuracy was not at the moment of grasp at time 0, but slightly earlier at −0.20*s*, 95%*CI* = [−0.32, −0.09]. This indicates that maximum information about the eye, head, and hand coordination is available just in time for the action.

**Figure 6:**
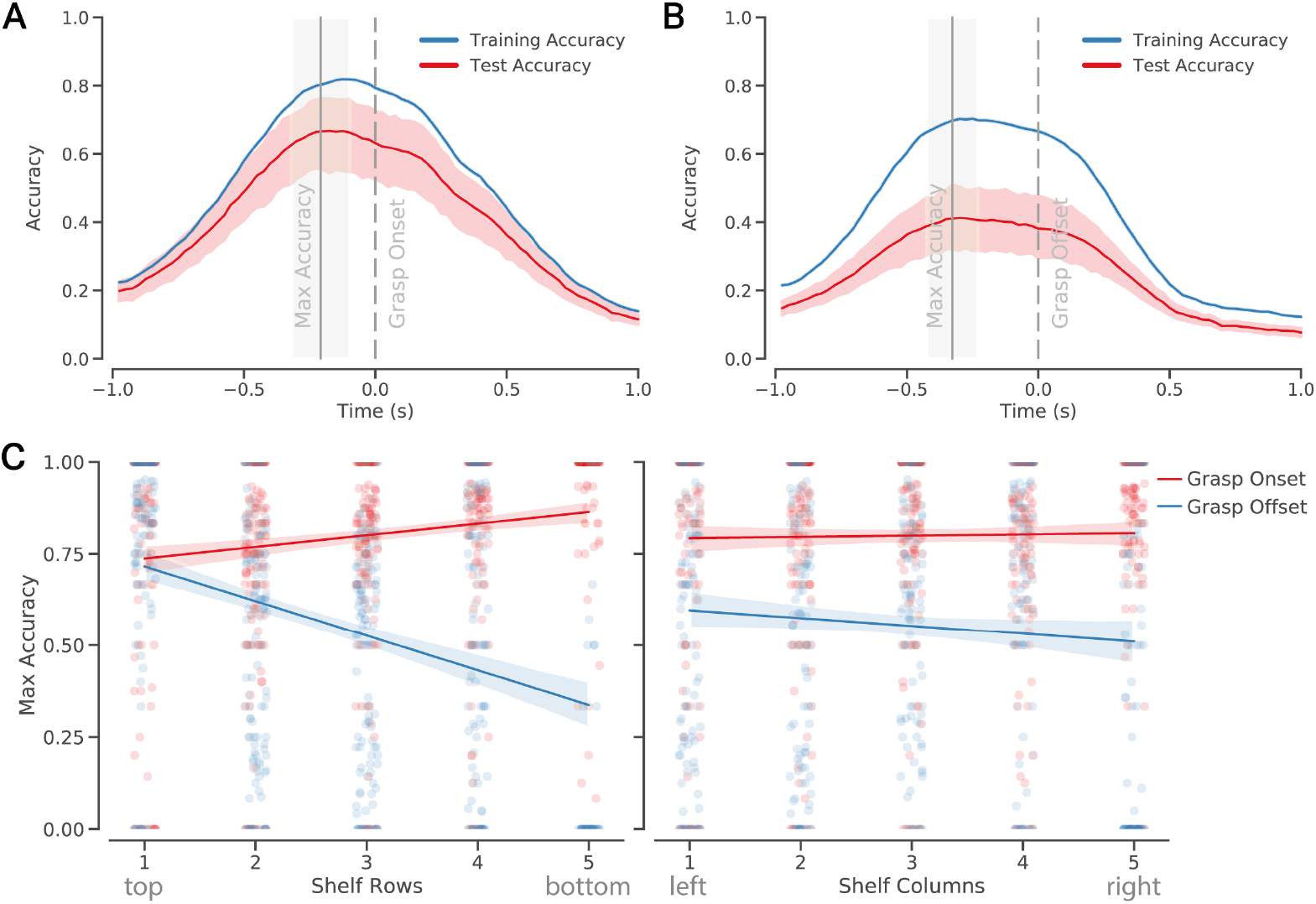
Accuracy of 2D PCA subspace in predicting the action location at each time point of the action epoch. **A**. The training (blue trace) and test/validation (red trace) accuracy for −1s before and 1s after the grasp onset events. Solid trace represents mean accuracy, and the shaded region denotes 95% CI of mean. **B**. The training and test accuracy for 1s before and 1s after grasp offset events. **C**. The linear fit over peak accuracy for the rows and columns of the predicted action location. Each dot represents the peak accuracy of a subject for grasp onset (red) and grasp offset(blue) events per row and column of the predicted action location on the shelf.

Similarly, the prediction of the object dropoff action locations across the grasp offset epoch is shown in Figure 6B. At 1s before the grasp offset, the predictive accuracy on the test data is 0.15, 95%*CI* = [0.13, 0.17]. At time point 0, the accuracy increases to 0.38, 95%*CI* = [0.30, 0.46] and decreases to 0.08, 95%*CI* = [0.06, 0.09] at 1s after the grasp offset event. Again, the maximum predictive accuracy was at time point −0.32*s*, 95%*CI* = [−0.42*s*, −0.23*s*] before the grasp offset. This further indicates the directional components of the eye, head, and hand vectors are aligned for action just in time.

With the above analysis, we generalized the predictive accuracy of the 2D PCA subspace. We determined the predictive information encapsulated in the subspace and the timing of the best prediction. For both grasp onset and grasp offset, the PCA space’s predictive accuracy increased with the approaching event and decreased after. Moreover, maximum accuracy is achieved just-in-time of the action event. This can be accounted for by the large explained variance ratio of the 2D subspace and the corresponding high correlations between the eye, head, and hand orientation vectors at that moment.

### Sources of Variance in Prediction

During grasp onset and especially grasp offset events, the validation accuracy is lower than the training accuracy and shows high variance. We looked into the source of this variance by separating the prediction accuracy by the shelf location of the action. We used a linear model to test the hypothesis that the row and column location of the upcoming action affected the predictive accuracy of the PCA-transformed 3D movement vectors. As shown in Figure 6C, the maximum accuracy increased linearly from the top to the bottom row in grasp onset events (*β* = 0.02, 95%*CI* = [0.01, 0.03], *t*(633.12) = 4.55, *p* = < 0.001). The maximum accuracy was not significantly affected during grasping objects from left to right columns (*β* = 0.00001, 95%*CI* = [− 0.01, 0.01], *t*(632.91) = −0.02, *p* = 0.987). In the case of grasp off-set events, the peak accuracy decreased significantly from top to bottom rows (*β* = −0.11, 95%*CI* = [−0.13, −0.10], *t*(555.10) = −11.96, *p* = < 0.001). In contrast, the peak accuracy decreased significantly but with a smaller slope across the shelf columns (*β* = −0.03, 95%*CI* = [−0.05, −0.02], *t*(555.15) = −3.95, *p* = < 0.001). Hence, the predictive accuracy of the PCA-transformed features was highly dependent on where the action would happen. The results show that the accuracy for grasp onset events did not differ dramatically across the different rows and columns of the shelf. However, there was a pronounced decrease in accuracy for grasp offset events, especially for lower shelf regions.

The maximum accuracy shows that the best information to predict the upcoming action is available just-in-time. However, the quality of this signal is lower for grasp offset events and significantly decreases for drop-off actions on the lower and rightward areas of the shelf, indicating a low correlation between the effectors for these locations.

## DISCUSSION

Our study explored the low-dimensional representations of natural visuomotor coordination. Subjects exhibited complex translation and rotation movements with their eyes, head, and right hand by making reaching movements to pick up and place objects on a life-size shelf in VR. We applied a time-wise PCA on the position and orientation vectors of the different effectors to capture the explained variance at each time point relative to grasp onset and offset events. Our analysis showed the complex system composed of the eye, head, and hand could be well described in a 2D PCA subspace. The PCA subspace showed an increase in the explained variance ratio at grasping events (onset and offset), where more than 60% of the variance is accounted for by the first two eigenvectors. Our analysis demonstrates a dynamic and distinct coupling of the horizontal and vertical components of effectors just in time for the upcoming action. Furthermore, this coupling showed high predictive accuracy of the target location of the forthcoming action. However, the accuracy was substantially influenced by the horizontal and vertical target location on the shelf. Hence, our study demonstrates a dynamic coupling and decoupling of eye, head, and hand movement vectors that have distinct features in the horizontal and vertical axes as well as dependent on the location of the reach target location.

### Methodological Considerations

Eye-hand or eye-head coordination is usually studied under constrained settings. In most cases, the horizontal and vertical positions or directions of the eye, head, and hand are extracted and directly correlated. In natural behavior, the complexity of the system does not afford such simplistic measures, as variables can interact with each other in non-obvious ways. Our study explored the latent relationships between the eye, head, and hand orientations in a low-dimensional space, helping to understand the underlying structure and relationships in the data relative to the action-critical events. We aimed to capture both the explained variance and the evolution of the variable loadings across time.

Our analysis of the cosine similarity of the original variables in the PCA subspace revealed strong associations between the horizontal and vertical components of the eye, head, and hand vectors. PCA transforms variables into principal components based on their variances and covariances. When using cosine similarity to assess relationships between original variables based on their loadings, it’s crucial to note that PCA mainly focuses on explaining variance, not necessarily revealing direct correlations between variables. It’s important to clarify that cosine similarity measures the angle between two vectors and is a measure of orientation similarity rather than a direct measure of statistical correlation in the traditional sense (Pearson’s correlation). Cosine similarity is less sensitive to the magnitude of vectors and focuses on their direction, which could be a limitation when the scale or variability of the original variables is relevant to their interpretation. In order to avoid drawing improper inferences, we plotted the distribution of the variable loadings on the first two PCs. We can confirm that the horizontal and vertical component loadings of the eye, head and hand had similar magnitudes on the PCs. Hence, by comparing the directionality of loadings, we could identify which variables share similar directional influences on the PCs, indicating underlying correlations that are not immediately obvious from the PCA results alone.

Finally, given the present study’s naturalistic setting, various noise sources could affect the findings. The noise source could be the eye or body trackers, which could exhibit errors due to slippage [38] or calibration errors [39]. We calibrated the trackers after every three trials to mitigate such errors. Moreover, as participants performed the task while wearing the VR head-mounted display, the head movements could have been cumbersome when picking up objects from the lower shelf locations. We did not direct participants to move in any one particular manner and asked them to make movements that were comfortable for them. Nonetheless, the insights offered by our study open the door to further experimental replications to validate our findings.

### Independence of Movement Vectors in the Horizontal and Vertical Axes

The cosine similarity analysis showed that eye, head, and hand coordination has distinct correlations between horizontal and vertical axes. While the vertical components of the eye, head, and hand were highly correlated during the investigated time windows, the eye-head and eye-hand horizontal components were correlated close to the action onset but were otherwise uncorrelated. In non-human primates, there is evidence of distinct areas in the premotor neural circuits for independent generation and control of saccadic movements in the horizontal and vertical axes [40]. Similar to the saccadic system, horizontal and vertical head movements are controlled by distinct circuits in the cerebellum [41] and the brainstem [42]. Similarly, premotor areas, primary motor cortex, and parietal cortex are differentially tuned for direction, with some studies indicating specialized subpopulations for horizontal versus vertical hand movements [43, 44, 45, 46]. Hence, there is ample evidence from primate studies showing distinct circuits that operate in a coordinated manner to produce smooth, multi-directional movements. For example, horizontal and vertical control centers are simultaneously active when making oblique hand or head movements, allowing for complex movement patterns [47]. Thus, our study offers preliminary behavioral evidence of independent but coordinated eye, head, and hand movement control in the horizontal and vertical axes in humans.

### Synergistic Coupling of the Effectors

The vertical components of the eye, head, and hand varied in the same direction and contributed significantly to the overall variance explained. Conversely, the horizontal component of the eye direction vector aligned with the head and hand horizontal components shortly before the action onset. Previous studies have shown that head and eye movements are generated by simultaneously receiving the same motor commands [48]. Here, head movements are necessary to center gaze in the orbits. Our data implies that head movements facilitate the vertical directionality of the eye, and the eye completes the last leg of the operation by making horizontal adjustments. Hence, the vertical components of the eye and head direction vectors were highly correlated, whereas the horizontal components showed a degree of independence.

The horizontal and vertical components of the head and hand were aligned throughout the grasping epoch. Studies have corroborated this strong coupling between the head and arm with respect to the eye in unrestrained macaque monkeys [16] and humans [15]. These studies argue that head movements facilitate foveation on the target to guide the final stages of object manipulation, leading to large correlations between the head and hand movements. In a similar vein, [17] showed a strong linkage between the head and hand movement trajectories, while the eye has a synergistic relationship instead of an obligatory one with them. Possibly this strong head-arm coupling results from learned motor behaviors during feeding where the head and hand orientations are coordinated to bring food to the mouth [49]. Head and hand coordination is not commonly studied, our results suggest the strong coupling between the two must be a consequence of a common neural code that drive this behavior.

### Differences between Grasp Onset and Grasp Offset

To validate the generalization of the PCA subspace, we predicted the grasp onset and offset locations by transforming the test data with the PCA weights of the training data. Since we cross-validated the model with unseen data, the test accuracy was expected to be lower than the training accuracy. During the reaching-to-grasp time window, the prediction accuracy across time on the test data is similar to the training data. However, the prediction accuracy on the test data is considerably lower for grasp offset events. This indicates idiosyncratic coordination to guide objects and drop them to desired locations, leading to lower explained variance and low correlation. Consequently, the 2D PCA subspace is likely less informative for grasp offset events.

Furthermore, the predictive accuracy had a larger variance for grasp offset events. Upon inspection, we found that the source of this variance is the location of the upcoming actions, where the accuracy decreased linearly from top to bottom rows and from left to right columns. The decreased accuracy further indicates that eye, head, and hand coordination is noisy and less informative in predicting upcoming actions. There is some evidence showing that eye-hand coordination is less precise when dropping off objects than when picking up [50, 51, 52]. Visual feedback is essential for adjusting the hand position in response to object features, so eye-hand coordination is particularly strong during grasping. Although visual feedback is necessary during object drop-off, haptic and proprioceptive feedback play a more dominant role, leading to less overt object-tracking by the eyes. Also, during grasping, the gaze moves ahead of the hand before the hand makes contact with the target object. In contrast, during object release, the eye tends to move to the next task or location before the object is fully dropped off [2], indicating a lower reliance on real-time visual feedback during object release. Hence, the lower predictive accuracy of the movement vectors during grasp offset events and for particular shelf locations is likely due to less coordination between the effectors and a greater reliance on proprioceptive inputs.

### Neural Correlates of Multi-Effector Coordination

The high correlation between different effectors in a natural context indicates a common neural code that mediates visuomotor coordination. The posterior parietal cortex (PPC) in macaque monkeys codes the reach target location with respect to both eye and hand [53], and the PPC achieves this transformation “by vectorially subtracting hand location from target location, with both locations represented in eye-centered coordinates.” Neurons in the parietal area V6A also encode visuospatial and motor-related information in mixed body/hand-centered coordinates, including depth (alongside horizontal and vertical axes) instead of pure body or hand-centered coordinates [54]. This encoding allows for precise spatial representation essential for reaching and other goal-directed movements. Further, spatial locations are encoded flexibly and idiosyncratically across the macaque parietal cortex [55] supporting a variety of task demands. Similarly, in humans, there is growing consensus that the intraparietal sulcus (IPS) and anterior intraparietal area (AIP) are the at the core of visually guided reach and grasp movements [56]. Furthermore, the frontal eye fields (FEF), aside from controlling gaze shifts [57], may have a role in independent head control and is located adjacent to the dorsal premotor cortex (PMd), where neurons associated with both oculomotor and hand movement activities have been identified [58]. Thus, visuomotor coordination is achieved by a complex interplay between several brain regions, and there is mixed evidence of the reference frames utilized to accomplish this. Our study invites further research into this dynamic neural encoding of multi-effector coordination.

## CONCLUSION

We studied the low-dimensional representations of visuomotor coordination in natural behavior. The multi-dimensional data comprising of eye, head, and hand movement vectors could be decomposed to 2D representations that explained about 65% of the data’s variance. A closer look at the subspace structure showed that the head and hand movements had a strong positive correlation and contributed substantially to the explained variance across time. However, the eye-head and eye-hand movements had distinct correlations in the horizontal and vertical axes, with maximum correlation just-in-time of the action. These results show separate mechanisms of coordination where the head and hand are coordinated simultaneously, and the eye is coordinated synergistically for goal-oriented behavior.

## AUTHOR CONTRIBUTIONS

AK, PK, TS: conceived and designed the study. TS, PK: Procurement of funding. AK: data collection. AK, FB: data pre-processing. AK: data analysis. AK, MAW: initial draft of the manuscript. AK, MAW, TS, PK: revision and finalizing the manuscript. All authors contributed to the article and approved the submitted version. We would like to thank Shadi Derakhshan and Imke Mayer for helping with the data collection.

## DATA AVAILABILITY

The experimental data and analysis code can be found at https://osf.io/9edby/.

## SUPPLEMENTARY MATERIAL

Supplementary Movie 1 can be found at https://osf.io/7nwh2

### GRANTS

We are grateful for the financial support by the German Federal Ministry of Education and Research for the project ErgoVR (Entwicklung eines Ergonomie-Analyse-Tools in der virtuellen Realität zur Planung von Arbeitsplätzen in der industriellen Fertigung)-16SV8052. The funders had no role in study design, data collection and analysis, decision to publish, or preparation of the manuscript.

## Notes

### Competing Interest Statement

The authors have declared no competing interest.

### Summary of Updates

text changes based on reviewer comments

## REFERENCES

[1] Hayhoe, MM, Shrivastava, A, Mruczek, R, and Pelz, JB. “Visual memory and motor planning in a natural task”. en. In: Journal of vision 3.1 (2003), pp. 49–63. DOI: 10.1167/3.1.6.

[2] Land, M, Mennie, N, and Rusted, J. “The roles of vision and eye movements in the control of activities of daily living”. en. In: Perception 28.11 (1999), pp. 1311–1328. DOI: 10.1068/p2935.

[3] Hu, Y and Goodale, MA. “Grasping after a delay shifts size-scaling from absolute to relative metrics”. en. In: Journal of cognitive neuroscience 12.5 (Sept. 2000), pp. 856–868. DOI: 10.1162/089892900562462.

[4] Droll, JA and Hayhoe, MM. “Trade-offs between gaze and working memory use”. en. In: Journal of experimental psychology. Human perception and performance 33.6 (Dec. 2007), pp. 1352–1365. DOI: 10.1037/0096-1523.33.6.1352.

[5] Flanagan, JR, Bowman, MC, and Johansson, RS. “Control strategies in object manipulation tasks”. en. In: Current opinion in neurobiology 16.6 (Dec. 2006), pp. 650–659. DOI: 10.1016/j.conb.2006.10.005.

[6] Belardinelli, A, Stepper, MY, and Butz, MV. “It’s in the eyes: Planning precise manual actions before execution”. en. In: Journal of vision 16.1 (2016), p. 18. DOI: 10.1167/16.1.18.

[7] Johansson, RS, Westling, G, Bäckström, A, and Flanagan, JR. “Eye-hand coordination in object manipulation”. en. In: The Journal of neuroscience: the official journal of the Society for Neuroscience 21.17 (Sept. 2001), pp. 6917–6932. DOI: 10.1523/jneurosci.21-17-06917.2001.

[8] Land, MF and Furneaux, S. “The knowledge base of the oculomotor system”. en. In: Philosophical transactions of the Royal Society of London. Series B, Biological sciences 352.1358 (Aug. 1997), pp. 1231–1239. DOI: 10.1098/rstb.1997.0105.

[9] Land, MF and Hayhoe, M. “In what ways do eye movements contribute to everyday activities?” en. In: Vision research 41.25-26 (2001), pp. 3559–3565. DOI: 10.1016/s0042-6989(01)00102-x.

[10] Keshava, A, Nezami, FN, Neumann, H, Izdebski, K, Schüler, T, and König, P. “Just-in-time: Gaze guidance in natural behavior”. en. In: PLoS computational biology 20.10 (Oct. 2024), e1012529. DOI: 10.1371/journal.pcbi.1012529.

[11] Bowman, MC, Johansson, RS, and Flanagan, JR. “Eye-hand coordination in a sequential target contact task”. en. In: Experimental brain research. Experimentelle Hirnforschung. Experimentation cerebrale 195.2 (May 2009), pp. 273–283. DOI: 10.1007/s00221-009-1781-x.

[12] Danion, FR, Mathew, J, Gouirand, N, and Brenner, E. “More precise tracking of horizontal than vertical target motion with both the eyes and hand”. en. In: Cortex; a journal devoted to the study of the nervous system and behavior 134 (Jan. 2021), pp. 30–42. DOI: 10.1016/j.cortex.2020.10.001.

[13] Ballard, DH, Hayhoe, MM, and Pelz, JB. “Memory representations in natural tasks”. en. In: Journal of cognitive neuroscience 7.1 (1995), pp. 66–80. DOI: 10.1162/jocn.1995.7.1.66.

[14] Biguer, B, Jeannerod, M, and Prablanc, C. “The role of position of gaze in movement accuracy”. In: Attention and performance XI (1985), pp. 407–424.

[15] Smeets, JB, Hayhoe, MM, and Ballard, DH. “Goal-directed arm movements change eye-head coordination”. en. In: Experimental brain research. Experimentelle Hirnforschung. Experimentation cerebrale 109.3 (June 1996), pp. 434–440. DOI: 10.1007/BF00229627.

[16] Arora, HK, Bharmauria, V, Yan, X, Sun, S, Wang, H, and Crawford, JD. “Eye-head-hand coordination during visually guided reaches in head-unrestrained macaques”. en. In: Journal of neurophysiology 122.5 (Nov. 2019), pp. 1946–1961. DOI: 10.1152/jn.00072.2019.

[17] Pelz, J, Hayhoe, M, and Loeber, R. “The coordination of eye, head, and hand movements in a natural task”. en. In: Experimental brain research. Experimentelle Hirnforschung. Experimentation cerebrale 139.3 (Aug. 2001), pp. 266–277. DOI: 10.1007/s002210100745.

[18] Hessels, RS et al. “Gaze-action coupling, gaze-gesture coupling, and exogenous attraction of gaze in dyadic interactions”. en. In: Attention, perception & psychophysics 86.8 (Nov. 2024), pp. 2761–2777. DOI: 10.3758/s13414-024-02978-4.

[19] Stamenkovic, A, Stapley, PJ, Robins, R, and Hollands, MA. “Do postural constraints affect eye, head, and arm coordination?” en. In: Journal of neurophysiology 120.4 (Oct. 2018), pp. 2066–2082. DOI: 10.1152/jn.00200.2018.

[20] König, P, Melnik, A, Goeke, C, Gert, AL, König, SU, and Kietzmann, TC. “Embodied cognition”. In: 2018 6th International Conference on Brain-Computer Interface (BCI). ieeexplore.ieee.org, Jan. 2018, pp. 1–4. DOI: 10.1109/IWW-BCI.2018.8311486.

[21] Pennartz, CMA. “Consciousness, representation, action: The importance of being goal-directed”. en. In: Trends in cognitive sciences 22.2 (Feb. 2018), pp. 137–153. DOI: 10.1016/j.tics.2017.10.006.

[22] Ingram, JN and Wolpert, DM. “Naturalistic approaches to sensorimotor control”. en. In: Progress in brain research 191 (2011), pp. 3–29. DOI: 10.1016/B978-0-444-53752-2.00016-3.

[23] Hayhoe, M and Ballard, D. “Modeling task control of eye movements”. en. In: Current biology: CB 24.13 (July 2014), R622–8. DOI: 10.1016/j.cub.2014.05.020.

[24] Ballard, DH and Hayhoe, MM. “Modelling the role of task in the control of gaze”. en. In: Visual cognition 17.6-7 (Aug. 2009), pp. 1185–1204. DOI: 10.1080/13506280902978477.

[25] Ballard, DH, Hayhoe, MM, Li, F, and Whitehead, SD. “Hand-eye coordination during sequential tasks”. en. In: Philosophical transactions of the Royal Society of London. Series B, Biological sciences 337.1281 (Sept. 1992), 331–8, discussion 338–9. DOI: 10.1098/rstb.1992.0111.

[26] Keshava, A, Aumeistere, A, Izdebski, K, and Konig, P. “Decoding Task From Oculomotor Behavior In Virtual Reality”. In: ACM Symposium on Eye Tracking Research and Applications. ETRA ‘20 Short Papers Article 30. New York, NY, USA: Association for Computing Machinery, June 2020, pp. 1–5. DOI: 10.1145/3379156.3391338.

[27] Keshava, A, Gottschewsky, N, Balle, S, Nezami, FN, Schüler, T, and König, P. “Action affordance affects proximal and distal goal-oriented planning”. en. In: The European journal of neuroscience 57.9 (May 2023), pp. 1546–1560. DOI: 10.1111/ejn.15963.

[28] König, SU, Keshava, A, Clay, V, Rittershofer, K, Kuske, N, and König, P. “Embodied Spatial Knowledge Acquisition in Immersive Virtual Reality: Comparison to Map Exploration”. In: Frontiers in Virtual Reality 2 (2021). DOI: 10.3389/frvir.2021.625548.

[29] Nolte, D, Vidal De Palol, M, Keshava, A, Madrid-Carvajal, J, Gert, AL, Butler, EM von, Kömürlüoėlu, P, and König, P. “Combining EEG and eye-tracking in virtual reality: Obtaining fixationonset event-related potentials and event-related spectral perturbations”. en. In: Attention, perception & psychophysics (July 2024). DOI: 10.3758/s13414-024-02917-3.

[30] Mobbs, D, Wise, T, Suthana, N, Guzmán, N, Kriegeskorte, N, and Leibo, JZ. “Promises and challenges of human computational ethology”. en. In: Neuron 109.14 (July 2021), pp. 2224–2238. DOI: 10.1016/j.neuron.2021.05.021.

[31] Bialek, W. “On the dimensionality of behavior”. In: Proceedings of the National Academy of Sciences 119.18 (2022), e2021860119. DOI: 10.1073/pnas.2021860119. eprint: https://www.pnas.org/doi/pdf/10.1073/pnas.2021860119.

[32] Santello, M, Flanders, M, and Soechting, JF. “Postural hand synergies for tool use”. en. In: The Journal of neuroscience: the official journal of the Society for Neuroscience 18.23 (Dec. 1998), pp. 10105– 10115. DOI: 10.1523/JNEUROSCI.18-23-10105.1998.

[33] Sanger, TD. “Human arm movements described by a low-dimensional superposition of principal components”. en. In: The Journal of neuroscience: the official journal of the Society for Neuroscience 20.3 (Feb. 2000), pp. 1066–1072.

[34] Corbeil, RR and Searle, SR. “Restricted Maximum Likelihood (REML) Estimation of Variance Components in the Mixed Model”. In: Technometrics: a journal of statistics for the physical, chemical, and engineering sciences 18.1 (Feb. 1976), pp. 31–38. DOI: 10.1080/00401706.1976.10489397.

[35] Luke, SG. “Evaluating significance in linear mixed-effects models in R”. en. In: Behavior research methods 49.4 (Aug. 2017), pp. 1494–1502. DOI: 10.3758/s13428-016-0809-y.

[36] Wilkinson, GN and Rogers, CE. “Symbolic Description of Factorial Models for Analysis of Variance”. In: Journal of the Royal Statistical Society. Series C, Applied statistics 22.3 (1973), pp. 392– 399. DOI: 10.2307/2346786.

[37] Sailer, U, Flanagan, JR, and Johansson, RS. “Eye-hand coordination during learning of a novel visuomotor task”. en. In: The Journal of neuroscience: the official journal of the Society for Neuroscience 25.39 (Sept. 2005), pp. 8833–8842. DOI: 10.1523/JNEUROSCI.2658-05.2005.

[38] Niehorster, DC, Santini, T, Hessels, RS, Hooge, ITC, Kasneci, E, and Nyström, M. “The impact of slippage on the data quality of head-worn eye trackers”. en. In: Behavior research methods 52.3 (June 2020), pp. 1140–1160. DOI: 10.3758/s13428-019-01307-0.

[39] Ehinger, BV, Groß, K, Ibs, I, and König, P. “A new comprehensive eye-tracking test battery concurrently evaluating the Pupil Labs glasses and the EyeLink 1000”. en. In: PeerJ 7 (July 2019), e7086. DOI: 10.7717/peerj.7086.

[40] Moschovakis, AK, Scudder, CA, and Highstein, SM. “The microscopic anatomy and physiology of the mammalian saccadic system”. en. In: Progress in neurobiology 50.2-3 (Oct. 1996), pp. 133–254. DOI: 10.1016/s0301-0082(96)00034-2.

[41] Shaikh, AG, Meng, H, and Angelaki, DE. “Multiple reference frames for motion in the primate cerebellum”. en. In: The Journal of neuroscience: the official journal of the Society for Neuroscience 24.19 (May 2004), pp. 4491–4497. DOI: 10.1523/JNEUROSCI.0109-04.2004.

[42] Crawford, JD, Henriques, DYP, Medendorp, WP, and Khan, AZ. “Ocular kinematics and eye-hand coordination”. en. In: Strabismus 11.1 (Mar. 2003), pp. 33–47. DOI: 10.1076/stra.11.1.33.14094.

[43] Georgopoulos, AP, Kalaska, JF, Caminiti, R, and Massey, JT. “On the relations between the direction of two-dimensional arm movements and cell discharge in primate motor cortex”. en. In: The Journal of neuroscience: the official journal of the Society for Neuroscience 2.11 (Nov. 1982), pp. 1527–1537. DOI: 10.1523/jneurosci.02-11-01527.1982.

[44] Kalaska, JF, Caminiti, R, and Georgopoulos, AP. “Cortical mechanisms related to the direction of two-dimensional arm movements: relations in parietal area 5 and comparison with motor cortex”. en. In: Experimental brain research 51.2 (1983), pp. 247–260. DOI: 10.1007/bf00237200.

[45] Hadjidimitrakis, K, De Vitis, M, Ghodrati, M, Filippini, M, and Fattori, P. “Anterior-posterior gradient in the integrated processing of forelimb movement direction and distance in macaque parietal cortex”. en. In: Cell reports 41.6 (Nov. 2022), p. 111608. DOI: 10.1016/j.celrep.2022.111608.

[46] Lacquaniti, F. “Representing spatial information for limb movement: Role of area in the”. In: Cerebral Cortex Scp/Oct 5 (1995), pp. 391–409.

[47] Crawford, JD, Henriques, DYP, and Medendorp, WP. “Three-dimensional transformations for goaldirected action”. en. In: Annual review of neuroscience 34.1 (2011), pp. 309–331. DOI: 10.1146/annurev-neuro-061010-113749.

[48] Land, MF. “Predictable eye-head coordination during driving”. en. In: Nature 359.6393 (Sept. 1992), pp. 318–320. DOI: 10.1038/359318a0.

[49] Hadjidimitrakis, K. “Coupling of head and hand movements during eye-head-hand coordination: there is more to reaching than meets eye”. en. In: Journal of neurophysiology 123.5 (May 2020), pp. 1579–1582. DOI: 10.1152/jn.00099.2020.

[50] Johansson, RS and Flanagan, JR. “Coding and use of tactile signals from the fingertips in object manipulation tasks”. en. In: Nature reviews. Neuroscience 10.5 (May 2009), pp. 345–359. DOI: 10.1038/nrn2621.

[51] Hesse, C and Deubel, H. “Efficient grasping requires attentional resources”. en. In: Vision research 51.11 (June 2011), pp. 1223–1231. DOI: 10.1016/j.visres.2011.03.014.

[52] Smeets, JB and Brenner, E. “A new view on grasping”. en. In: Motor control 3.3 (July 1999), pp. 237– 271. DOI: 10.1123/mcj.3.3.237.

[53] Buneo, CA, Jarvis, MR, Batista, AP, and Andersen, RA. “Direct visuomotor transformations for reaching”. en. In: Nature 416.6881 (Apr. 2002), pp. 632–636. DOI: 10.1038/416632a.

[54] Hadjidimitrakis, K, Ghodrati, M, Breveglieri, R, Rosa, MGP, and Fattori, P. “Neural coding of action in three dimensions: Task- and time-invariant reference frames for visuospatial and motor-related activity in parietal area V6A”. en. In: The Journal of comparative neurology 528.17 (Dec. 2020), pp. 3108–3122. DOI: 10.1002/cne.24889.

[55] Chang, SWC and Snyder, LH. “Idiosyncratic and systematic aspects of spatial representations in the macaque parietal cortex”. en. In: Proceedings of the National Academy of Sciences of the United States of America 107.17 (Apr. 2010), pp. 7951–7956. DOI: 10.1073/pnas.0913209107.

[56] Culham, JC, Cavina-Pratesi, C, and Singhal, A. “The role of parietal cortex in visuomotor control: what have we learned from neuroimaging?” en. In: Neuropsychologia 44.13 (2006), pp. 2668–2684. DOI: 10.1016/j.neuropsychologia.2005.11.003.

[57] Tu, TA and Keating, EG. “Electrical stimulation of the frontal eye field in a monkey produces combined eye and head movements”. en. In: Journal of neurophysiology 84.2 (Aug. 2000), pp. 1103– 1106. DOI: 10.1152/jn.2000.84.2.1103.

[58] Fujii, N, Mushiake, H, and Tanji, J. “Rostrocaudal distinction of the dorsal premotor area based on oculomotor involvement”. en. In: Journal of neurophysiology 83.3 (Mar. 2000), pp. 1764–1769. DOI: 10.1152/jn.2000.83.3.1764.

